# Mass-planted large bromeliads in an urban landscape increase the risk for mosquitoes of pest and public health concern

**DOI:** 10.1101/2025.10.06.680812

**Authors:** Pui Yin A. Lok, Victoria J. Brookes, Cameron Webb

**Affiliations:** Sydney School of Veterinary Science, Faculty of Science, The University of Sydney, Camperdown, New South Wales, 2006, Australia; Sydney Infectious Diseases Institute, Faculty of Medicine and Health, The University of Sydney, Camperdown, New South Wales, 2006 Australia; School of Medical Sciences, Faculty of Medicine and Health, The University of Sydney, Camperdown, New South Wales, 2006 Australia; Department of Medical Entomology, NSW Health Pathology, Westmead Hospital, Westmead, New South Wales, 2145 Australia

**Keywords:** Australia, Bromeliaceae, Arbovirus, Mosquito-borne diseases, Sydney, Urban greening

## Abstract

Exotic bromeliads (Bromeliaceae) are increasingly used in Australian urban green spaces for their hardiness and aesthetic appeal. However, the water-holding tanks and axils of these plants provide habitat for mosquitoes, raising public health concerns that must be balanced with ecological objectives of urban greening. This study investigated the abundance and species richness of immature and adult mosquitoes associated with bromeliad plantings in Sydney, Australia.

Between October 2023 and April 2024, immature mosquitoes were sampled weekly from large bromeliads at 17 locations, and adults were collected with CO-baited Encephalitic Virus Surveillance traps at six sites of contrasting bromeliad density. Specimens were identified to species level, and associations with climatic variables, bromeliad species, and planting characteristics were analysed using generalised linear mixed models.

A total of 2,326 immature mosquitoes (three species) and 6,366 adult mosquitoes (ten species) were collected. Aedes notoscriptus and Culex quinquefasciatus dominated both life stages. Total immature abundance increased by 4% for each additional bromeliad in a patch (IRR 1.04, 95% CI 1.02–1.05), and was highest in Alcantarea imperialis (IRR 1.31, 95% CI 0.97–1.77). Weekly-lagged humidity was positively associated with immature counts (IRR 1.01 per 1% increase, 95% CI 1.00–1.03). Adult abundance was significantly higher at high-density compared to low-density bromeliad sites (IRR 0.37, 95% CI 0.23–0.60).

Mass plantings of large, water-holding bromeliads might substantially increase mosquito populations in urban environments, elevating pest and public health risks. This highlights the need for integrated mosquito habitat management in sustainable landscaping design and planning.

**Highlights:**

- Mass bromeliad plantings could substantially increase urban mosquito populations and public health risks.
- Sites with high-density bromeliads yielded 170% more adult mosquitoes than low-density sites.
- *Aedes notoscriptus* and *Culex quinquefasciatus* dominated both immature and adult mosquito collections.
- Findings inform sustainable urban landscaping strategies to reduce the pest and public health risks associated with mosquitoes.

## 1 Introduction

Over recent decades, global anthropogenic climate changes and increased urbanization have increased the interface between humans, animals, and ecosystem environments, potentially leading to higher transmission of both human and animal diseases (Andersen & Davis, 2017; Perrin et al., 2022; Vora, 2008). Mosquitoes (Diptera: Culicidae) are widespread in urban centers around the world and many species are vectors of the pathogens that cause mosquito-borne diseases (MBDs), including both parasites and arboviruses (World Health Organisation, 2024). The mortality, morbidity, and economic burden of mosquito-borne diseases globally is significant, with >700 million cases of disease, >1 million deaths, and US $12 billion burden annually (Andersen & Davis, 2017; Caraballo & King, 2014; Chilakam et al., 2023; World Health Organisation, 2024).

Mosquitoes generally have a short but complex life cycle, with entirely aquatic immature stages (C. Webb et al., 2016a) and they have adapted to most water-containing ecological niches and climate zones. (Hall & Tamïr, 2022; C. Webb et al., 2016a). While most mosquito species show a preference for specific types of egg-laying habitats and have high sensitivity to environmental changes, some are more resilient and tend to increase in human-modified environments, such as urban landscapes (Dorvillé, 1996; Juliano & Lounibos, 2005). Such landscapes favour several vector mosquitoes, including *Aedes aegypti* and *Culex quinquefasciatus,* which demonstrate preferences for natural or artificial water-holding containers or polluted waterbodies for egg-laying (Medeiros-Sousa et al., 2017).

Plants from the family Bromeliaceae are a natural water-holding container that can provide a suitable egg-laying habitat for these mosquitoes (Docile et al., 2017; Mocellin et al., 2009; Wilke et al., 2018). They are a family of monocotyledonous plants native to the Neotropics, with the extension of a few species into West Africa. They are classified into three subfamilies (Pitcairnioideae, Bromelioideae, and Tillandsioideae), consisting of >70 genera and >2500 species (Holst & Luther, 2004). Bromeliads have an inferior ovary encircled by closely overlapping central leaves which can retain rainwater and organic debris in the leaves and stems to acquire nutrients (Picado, 1913). This central well and the four to six extra pockets created by the outer axial leaves serve as a habitat for numerous terrestrial arthropods to undergo aquatic immature stages, such as mosquitoes, non-biting midges (Diptera: *Chironomidae*), and moth flies (Diptera: *Psychodidae*) (Frank & Lounibos, 2009; Ladino et al., 2019). Studies in the United States and Brazil found immature mosquitoes of three genera (*Culex*, *Aedes*, and *Wyeomyia*) in native bromeliads planted in residential areas (Mocellin et al., 2009; Wilke et al., 2018). There is also the suggestion that the international trade in bromeliads may be contributing to the spread of exotic mosquitoes (Multini et al., 2020; Vajda et al., 2018). Therefore, bromeliads could serve as habitat for many mosquito species, which could have health implications from MBDs in urban areas, especially given their increased popularity in plantings of urban green spaces due to their attractiveness and hardiness (Marques et al., 2012; Mocellin et al., 2009).

As urbanization increases, global efforts aim to reduce its negative environmental and social impacts by implementing strategies, such as sustainable landscaping and ecosystem conservation (R. Karade et al., 2017; Koh & Sodhi, 2004). Sustainable landscaping is an approach that aims to limit negative environmental impacts and reduce pollution (Loehrlein, 2020) through the creation of urban green space to enhance ecosystem service (Peng et al., 2017; Pham et al., 2022). Sydney, Australia, is a major metropolitan region at risk of adverse impacts of heatwaves and extreme weather events linked to changing climate (Chaston et al., 2022). There is an increasing focus on ‘green infrastructure’ in urban areas to mitigate the impact of heat (Bartesaghi-Koc et al., 2020) and in Sydney, urban green space covers around 57%, with >400 parks containing diverse native and exotic flora (Hsu et al., 2022).

Bromeliads are commonly used in these urban green spaces due to their colourful appearance and tolerance of hot and dry conditions, greatly reducing the burden of maintenance (Shultis, 2009). However, this use of bromeliads in urban landscapes potentially provides habitats for mosquitoes (Marques et al., 2012). Many endemic and exotic mosquitoes in Australia have been shown to be associated with bromeliads, such as *Aedes aegypti*, a known vector of dengue virus (DENV) in Central and Far North Queensland, as well as *Aedes notoscriptus* and *Culex quinquefasciatus,* two vectors of pathogens of human and veterinary health concern (C. Webb et al., 2016a). Historic populations of *Ae. aegypti* from Sydney have been found to breed in bromeliads under experimental conditions (Shultis, 2009) and populations elsewhere in Australia have been found in associations with bromeliads (Williams et al., 2013). A common and widespread pest and vector mosquito is *Ae. notoscriptus,* a known vector of Ross River virus (RRV) in Australia (Watson & Kay, 1998) and closely associated water-holding containers, including bromeliads, in urban areas. Surveys of residential properties in many parts of Australia have identified bromeliads as actual and potential habitat for this mosquito (Kay et al., 2008), there is a paucity of understanding of the relationship between bromeliads used in public space landscaping and immature mosquito dynamics in the Australian urban landscape.

Given that outbreaks of MBDs in some suburbs of metropolitan Sydney have been reported, gaps in understanding the ecological dynamics between mosquito vectors, bromeliads, and urban environments is essential for managing public health risks while pursuing sustainable landscaping. Understanding the relationship between bromeliad planting and mosquito dynamics is important for managing public health risks while pursuing sustainable landscaping (Amin et al., 1998; Brokenshire et al., 2000; C. Webb et al., 2001).

This study aimed to quantify mosquito abundance and species richness in bromeliad plantings in Sydney’s urban landscape and determine how bromeliad species and abundance, planting density, and environmental factors influence mosquito populations. We hypothesized that bromeliads serve as productive mosquito habitats that significantly contribute to local mosquito populations, with effects varying by plant species and planting configuration. Using the University of Sydney’s Camperdown campus as a model urban environment, we sampled both immature and adult mosquitoes to provide evidence-based guidance for sustainable urban landscaping practices that minimize mosquito-borne disease risks while maintaining ecological benefits.

## 2 Materials and methods

### 2.1 Study site

This observational study was conducted on the Camperdown Campus, The University of Sydney, Sydney, from October 2023 to April 2024 (7 months). Sydney has four distinct seasons: summer from December to February, autumn from March to May, winter from June to August, and spring from September to November (Bureau of Meterorology, n.d.). This site was selected due to an abundance of bromeliads as well as noteworthy variation in planting density throughout the site, and the sampling period was selected to coincide with the warmer seasons when mosquitoes are most active. The study was suspended from December 22, 2023, to January 1, 2024, and from March 4 to 16, 2024, due to the inaccessibility of the study area (seasonal holiday period and construction-related access restrictions). Camperdown is southwest to the Central Business District of Sydney, New South Wales, Australia (33°52’1.19” S, 151°12’16.20” E). This area is heavily urbanized, surrounded by medium to high-density residential development, educational institutions, and parks. The campus spans 72 hectares and features multiple buildings with gardens and lawns (https://www.sydney.edu.au/about-us/campuses/campus-locations.html, accessed 14 May 2025). These gardens are human-structured landscapes, primarily composed of a diverse flora of both native and exotic plants, such as native Sydney golden wattle (*Acacia longifolia*), small-leaved fig trees (*Ficus obliqua*), exotic Lion’s tail agaves (*Agave attenuata*), and bromeliads (*Bromeliaceae spp*.) (https://campusflora.sydney.edu.au/, accessed 14 May 2025). Many of these bromeliads are found in established garden beds under trees and in areas of shade.

### 2.2 Mapping of bromeliads distribution

The garden beds within the study site were surveyed for bromeliad plants in September 2023. Six species of bromeliad species (*Vriesea hieroglyphica, Alcantarea imperialis, Alcantarea nigripetala, Aechmea caudata, Aechmea fasciata,* and *Neoregelia marmorata*) (Fig. 1) were identified. Of these, the species that grows the largest is *A. imperialis* with a potential leaf height of up to 3m and a water capacity of up to 30L (Silva et al., 2022). The bromeliads were not irrigated, only exposed to natural rainfall and were not treated with insecticide during the sampling period nor in the previous six months (*pers. comm*. The University of Sydney). The height of each bromeliad was measured from the soil line to the highest leaf tip to categorize bromeliads into “small bromeliad” and “large bromeliad” based also on adult plant size. “Small bromeliads” included *Neoregelia marmorata* and *Alcantarea sp.* <1m in height, and *Aechmea spp., and Vriesea hieroglyphica* <50cm. “Large bromeliads” included *Alcantarea sp.* >1m in height, and *Aechmea sp.,* and *Vriesea hieroglyphica* >50cm in height (Fig. 1). Bromeliad plants in the “small bromeliad” category were excluded from the study, because the water contained in the central well of these plants was not accessible and prone to evaporation, making them difficult to sample and contributing less to mosquito breeding habitats. Locations with large bromeliads are shown in Figures 1 and S1 (with grid).

**Figure 1.**
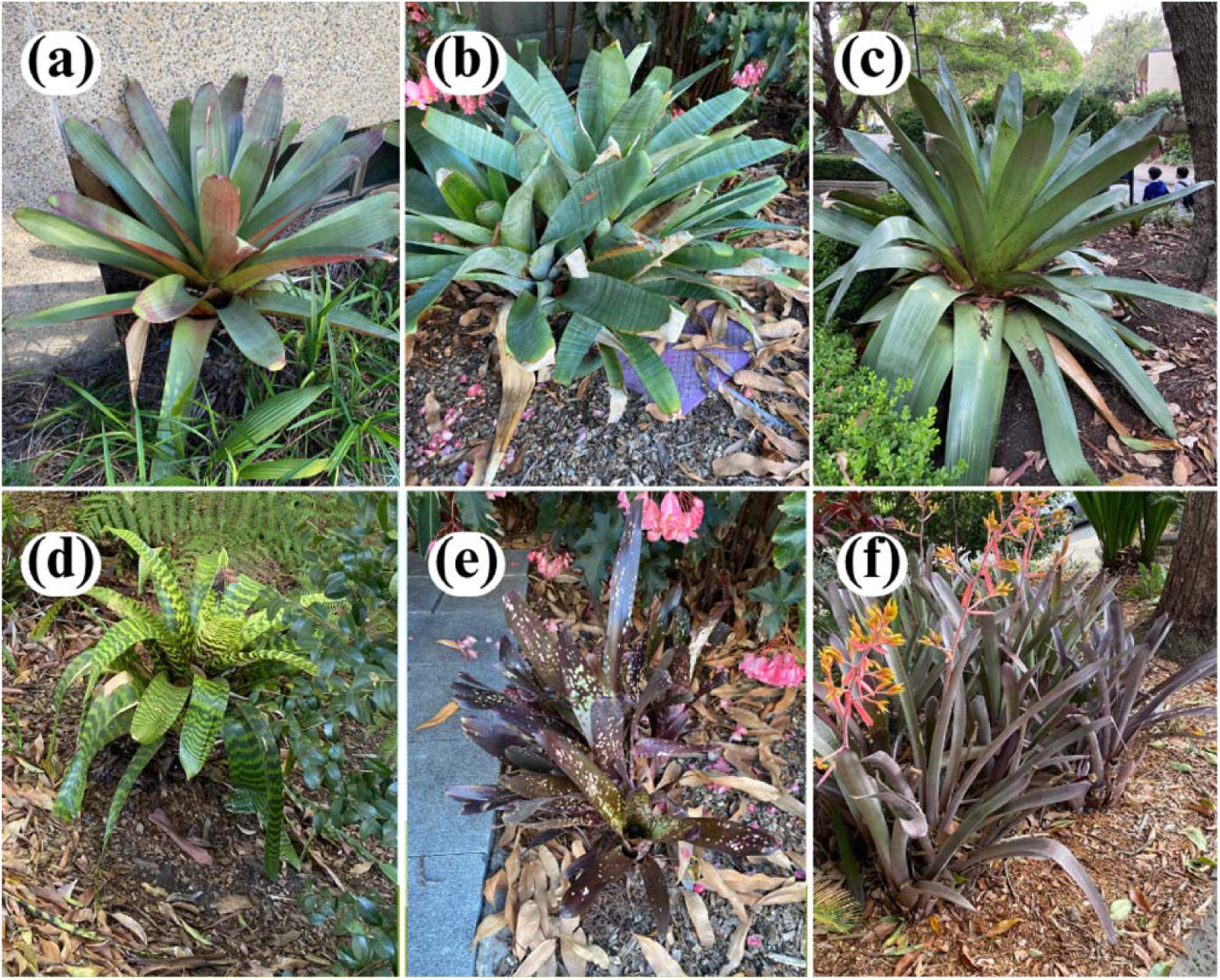
Bromeliad species at the Camperdown Campus, the University of Sydney. (a: *Alcantarea imperialis*; b: *Aechmea fasciata*; c: *Alcantarea nigripetala*; d: *Vriesea hieroglyphica*; e: *Neoregelia marmorata*; f: *Aechmea caudata*) *(Source: Pui Yin Lok)*.

**Figure 2.**
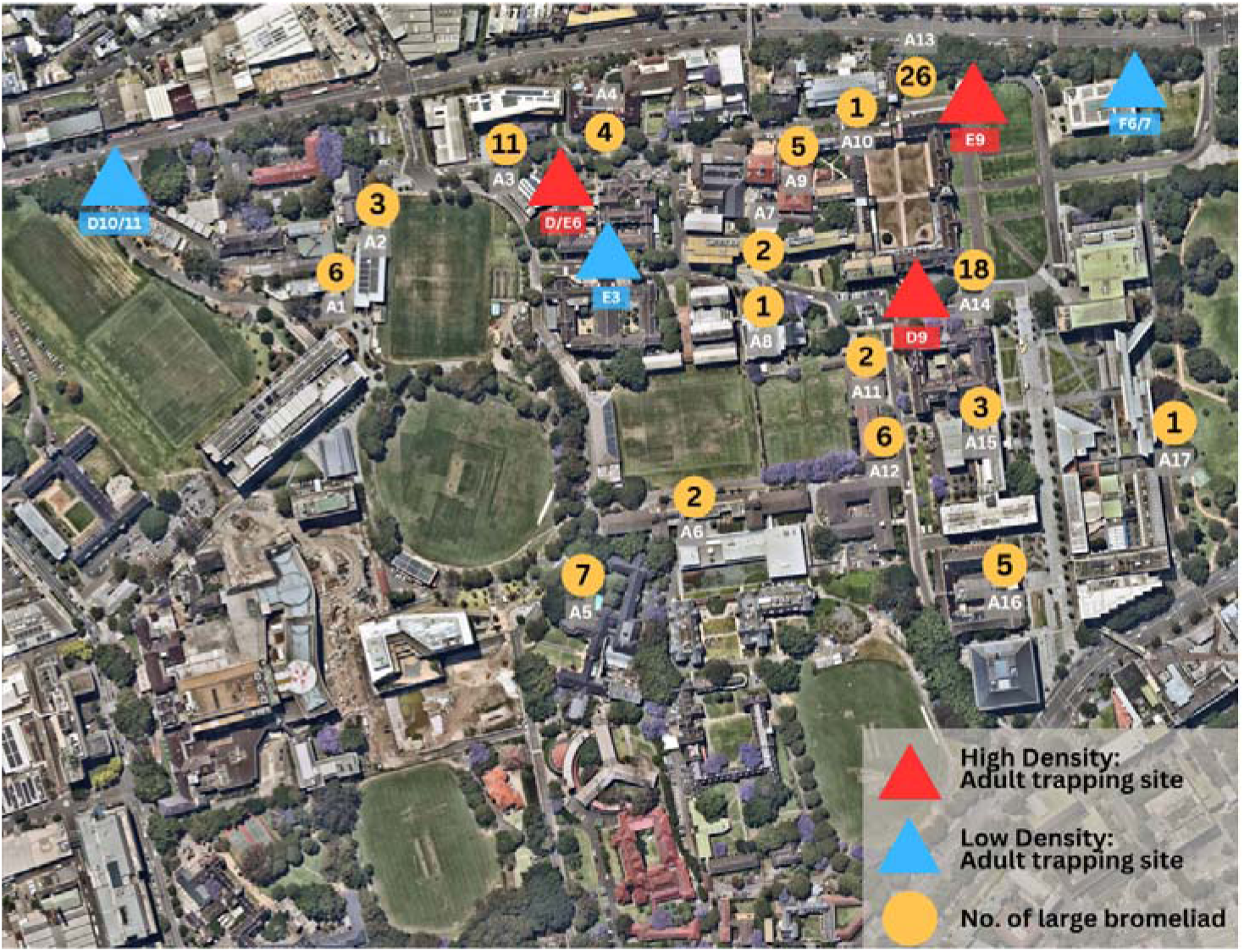
Study area showing 17 immature and 6 adult mosquito sampling locations in the Camperdown Campus, The University of Sydney to explore the abundance and species richness of mosquitoes between October 2023 and April 2024 (A1: R.M.C Gunn Building; A2: JD Stewart Building; A3: RD Watt Building; A4: Social Science Building; A5: St Paul’s College; A6: Physics Building; A7: Brennan MacCallum Building; A8: Manning House; A9: Pharmacy Building; A10: Botany lawn (Science Road); A11: Sidewalk of RC Mills Building; A12: RC Mills Building; A13: Botany lawn (Macleay Building); A14: Garden bed next to the Quadrangle; A15: Anderson Stuart Building; A16: Madsen Building; A17: Victoria Park; D10/11: Vet clinic; E3: Oval; F6/7: Museum; D/E6: Science Road west; D9: The Quadrangle; E9: Botany lawn)

The study site and surrounds were assessed for the presence of actual and potential non-bromeliad mosquito habitats. There were small numbers of artificial water-holding containers (for example, discarded plastic, glass, and metal objects) across the site that would be anticipated to be common in any urban landscape but were not considered likely to make a significant contribution to overall mosquito activity. There were two ornamental water features, but these had recirculated water and were not considered suitable mosquito habitat. Lake Northam (33°53′08″ S, 151°11′36″ E), a constructed pond within parklands adjacent to the Camperdown Campus was 200m from the closest adult mosquito trap and 230m from the closest large bromeliad.

### 2.3 Selection of immature and adult mosquito trapping sites

The study site was divided into 100x100m quadrats (Figure S1). A total of 14 quadrats contained “large bromeliads” (n = 103) in 17 distinct sites.

To determine sufficient sample size, a 1-stage freedom analysis was conducted using ‘Epitools’ (https://epitools.ausvet.com.au/, accessed 21 July 2025) to estimate the minimum number of bromeliads that needed to be sampled to detect a lowest expected immature mosquito prevalence (i.e. design prevalence). (Ausvet, n.d.). We assumed a diagnostic sensitivity and specificity of 100% for immature mosquito sampling, because all aquatic content from the central wall of each large bromeliad was aspirated, and identification of the presence or absence of immature mosquito is straightforward; therefore, there was a negligible probability of false positive and false negative samples. We set a design prevalence of 5% of large bromeliads containing immature mosquitoes if present, with a 95% confidence interval. Therefore, the required sample size for this study was 41 (40% of large bromeliads). Due to the ease of sampling, 50% of large bromeliads (n= 55) in each of the 17 locations were randomly selected for immature mosquito sampling (rounded up to the nearest whole number; for example, at sites with three bromeliads, two were sampled each week, and at sites with one bromeliad, the same bromeliad was sampled each week) each week. Randomization without replacement of bromeliads at each site was implemented using an online number generator (https://stattrek.com/statistics/random-number-generator, accessed 6^th^ October 2023) before each sampling session.

For adult mosquito sampling, six sites were chosen representing locations with low and high densities of “large bromeliads” (3 of each density level). To determine bromeliad density, a heatmap was produced using QGIS 3.36 software (https://qgis.org; accessed on 10 October 2023; Figure S1). High-density bromeliad sites (D/E6, D9, E9) were from randomly selected quadrats in the highest quartiles (>50^th^ percentile) for bromeliad density, while low-density bromeliad sites (D10/11, E3, F6/7) were selected from randomly selected quadrats in the lowest quartile for bromeliad density. Quadrats were then assessed for suitability of hanging adult mosquito traps, and if there were no suitable sites (for example, the square contained a building), another square was randomly selected until 3 locations in each density zone were selected.

### 2.4 Immature mosquito sampling

Immature mosquitoes were sampled weekly at each of the 17 sites from October 2023 to March 2024 using a 50 ml pipette (Australian Entomological Supplies, https://www.entosupplies.com.au/, accessed 3^rd^ April 2024) to aspirate all aquatic content from the central well of each large bromeliad. This method was selected because it has been validated by previous bromeliad ecology studies that this drains bromeliads’ water reservoir tanks (Marques et al., 2012; Wilke et al., 2018). Immature mosquitoes (including all stages of larvae and pupae) were removed and preserved in 70% ethanol for counting and identification under a stereo microscope using an existing taxonomic key (Russell et al., 1993). After the removal of immature mosquitoes, the remaining aquatic content was returned to the central wall of each bromeliad. Immature mosquitoes from all samples were counted and identified twice to minimize information bias.

### 2.5 Adult mosquito sampling

Adult mosquitoes were sampled four times (November, December 2023, January, and March 2024) at each of the six adult sampling sites using CO_2_ baited Encephalitic Virus Surveillance (EVS) traps (Australian Entomological Supplies, https://www.entosupplies.com.au/, accessed 3^rd^ April 2024). This trap type is commonly used for mosquito research and surveillance in Australia (Rohe et al., 1979) and has been validated to capture mosquitoes intact with minimal entomological bycatch (Roiz et al., 2012; Spitzen et al., 2008; Yi et al., 2014). The traps contained approximately 500g of dry ice (a source of carbon dioxide) as the primary attractant, and an incandescent light as the secondary attractant. Host-seeking mosquitos are blown into a container attached to the trap by a mesh sock that holds the mosquitoes until they are ready to be counted and identified.

At each of the six adult sampling sites, three fixed replicate trap sites were established at least 20m apart. Traps were placed in areas sheltered from direct sunlight and wind. Traps were deployed overnight (approximately 4pm - 8am). Mosquito collections were returned to the laboratory where the mosquito specimens were placed in a freezer (-20 °C) for 30 minutes to kill them before being stored in petri-dishes prior to identification. All mosquito specimens were identified to species under a stereo microscope using an existing taxonomic key (Russell et al., 1993) and pictorial guide (C. Webb et al., 2016b).

### 2.6 Weather data

Weekly precipitation, average wind speed, temperature, and humidity data were obtained from the Australian Bureau of Meteorology website (http://www.bom.gov.au/climate/data/, accessed 24^th^ March 2024), using records from the station closest to the site (Observatory Hill - station ID: 66214; Lat: 33.86° S, Lon: 151.20° E).

#### 2.7.1 Statistical Analysis

All data were entered in Microsoft Excel 2024 spreadsheets (Mustafy & Rahman, 2024). All statistical analyses were conducted using the R statistical program (The R Foundation, n.d.) with packages readxl (Wickham et al., 2019), lubridate (Grolemund & Wickham, 2011), dplyr (Yarberry, 2021), ggplot2 (Wickham, 2009), MASS (Ripley et al., 2013), pscl (Jackman et al., 2015), performance (Lüdecke, Ben-Shachar, et al., 2021), lme4 (De Boeck et al., 2011), and sjPlot (Lüdecke, Patil, et al., 2021). All statistical tests were conducted with a significance level of 5% (α = 0.05)

#### 2.7.2 Descriptive analyses

Descriptive statistics were generated for immature and adult mosquito abundance, species richness, and species composition throughout the study period. Species abundance refers to the number of individuals of each species, species richness refers to the total number of different species present, and species composition refers to the relative abundance (percentage) of each species within the mosquito community.

#### 2.7.2 Evaluation of environmental variables, sampling locations, and bromeliad species on the immature and adult mosquito abundance

The effects of weather variables (temperature, humidity, weekly rainfall, and average wind speed), bromeliad species, and patch size (number of bromeliads per location) on immature and adult mosquito abundance were assessed using generalized linear mixed models (GLMMs) with a Poisson or negative binomial distribution and log link as required to accommodate over- or under-dispersion (glmmTMB’ package in R; (Magnusson et al., 2017)). All covariates were included a-priori based on biological plausibility. Random intercepts were specified for bromeliad (immature-mosquito models) or trap identity (adult-mosquito models), and for sampling date, to account for spatial and temporal non-independence. Environmental predictors were lagged by one week to reflect the 5–7-day egg-to-pupae development period pupae (Lim et al., 2021; C. Webb et al., 2016a). Model selection was guided by the small-sample-corrected Akaike Information Criterion (AICc) (Burnham & Anderson, 2002), and diagnostics were performed using the ‘performance’(Lüdecke, Ben-Shachar, et al., 2021) and ‘DHARMa’ packages in-R (Hartig & Hartig, 2017), including tests for overdispersion, inspection of residual and random-effects patterns, and calculation of conditional R². Final outputs included fixed-effect estimates as incidence-rate ratios (exponentiated coefficients) with 95% CIs, and bias-corrected marginal predicted values (‘ggpredict’ function from the package ‘ggeffects’; (Lüdecke, 2018)) averaged over random-effect levels. Absolute (additive) effects were quantified as average marginal slopes on the response scale (marginaleffects package in R; (Arel-Bundock et al., 2024)).

## 3 Results

### 3.1 Bromeliad species richness and abundance

One hundred and three large bromeliads of three species – *V. hieroglyphica, A. imperialis,* and *A. nigripetala* (n = 2, 67, and 34, respectively) – were identified at the study site. Locations with greatest bromeliad abundance were Botany Lawn (Macleay Building) (A13) (15 *A. imperialis,* 11 *A. nigripetala*), garden bed next to the Quadrangle (A14) (18 *A. imperialis*), and R.D. Watt building (A3) (11 *A. nigripetala*). The J.D Stewart building (A2) was the only location with *V. hieroglyphica* (n = 2).

### 3.2 Immature mosquito species richness, abundance, and dynamics

From 6 October 2023 to 17 April 2024, all bromeliads were sampled at least once within the 25 survey days (total 1375 sampling events, median 13 samples/bromeliad, range 1-25 samples/bromeliad). Temperature varied from 17.2—38.7°C, humidity from 23—98%, rainfall 1.4—113.8mm, and wind speed from 4—76km/h.

From these samples, 2,326 immature mosquitoes were collected and identified, and comprised three species: *Ae. notoscriptus* (83.6%)*, Cx. quinquefasciatus*, and *Toxorhynchites speciosus* (Tables 1 and S1).

The total abundance of immature mosquitoes by location was strongly correlated with the number of bromeliads at each location, (Pearson’s ρ = 0.95 [95% confidence interval [CI] 0.87—0.98]). The weekly abundance of immature mosquitoes ranged 6—259 individuals and was lowest from 2 October to 6 November 2023 and highest from 2 –7 January 2024 (Figure 3). Median immature mosquito abundance per bromeliad on each sampling occasion was 0 (range: 0–25; Figure 4).

**Figure 3.**
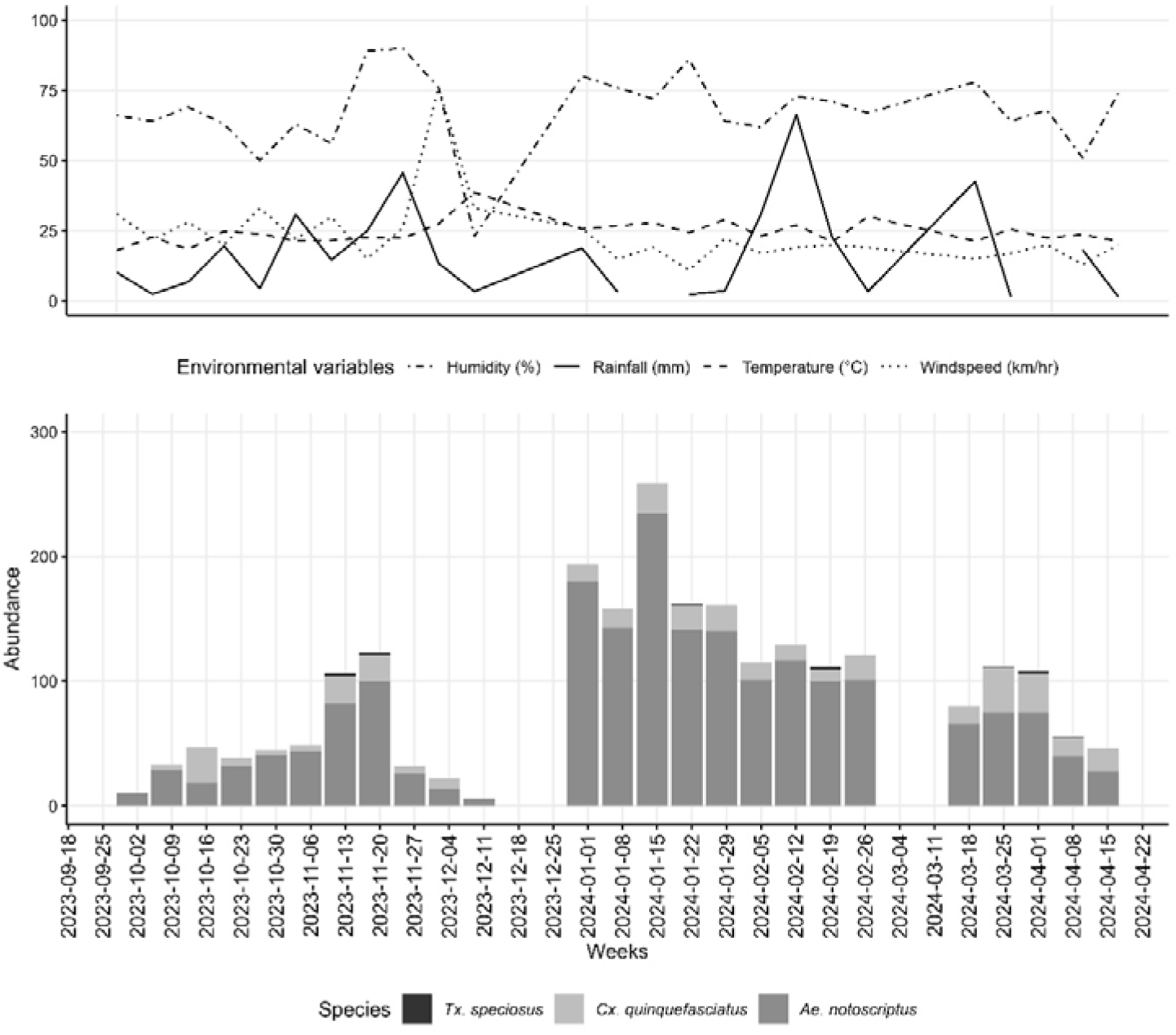
Weekly immature mosquito abundance in large bromeliads and weather variables at Camperdown campus, The University of Sydney, October 2023—April 2024.

**Figure 4.**
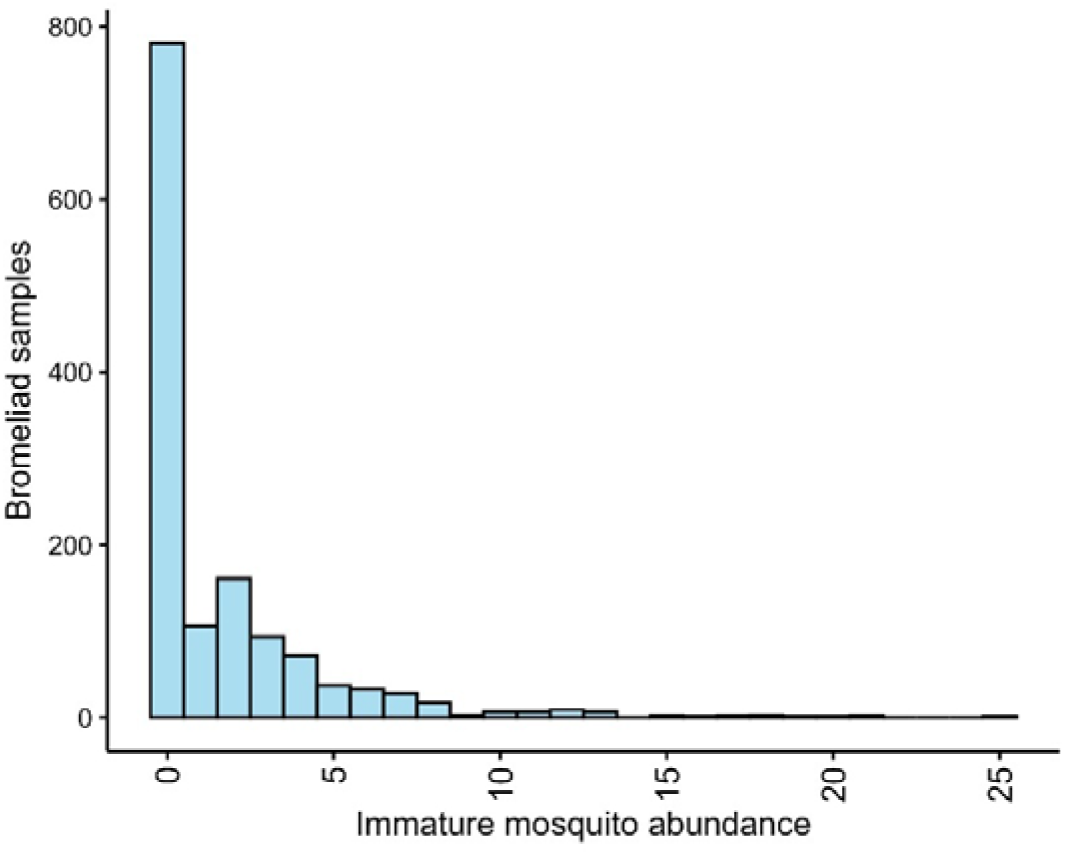
Histogram of immature mosquito abundance in large bromeliads at Camperdown campus, The University of Sydney October 2023—April 2024.

An initial Poisson GLMM for immature mosquito counts with one□week□lagged weather predictors, patch size, bromeliad species, and random intercepts for bromeliad identity (n = 103) and sampling date (n = 25), showed acceptable dispersion (dispersion ratio□=□0.73, *P*□=□0.94) but significant zero inflation (zero□inflation ratio□=□1.43, *P*□<□0.001). A zero□inflated Poisson GLMM (intercept□only zero□inflation component) avoided overdispersion (dispersion ratio□=□0.71, *P*□=□0.75) and zero inflation (ratio□=□0.99, *P*□=□0.99) with improved AICc (AICc□= 4406 *vs*. 5363), but a negative□binomial GLMM (same random□effect structure) did not have significant under-dispersion (dispersion□ratio =□0.35, *P*□=□0.1) and no zero inflation (ratio□=□1.04, *P*□=□0.49) with the lowest AICc (4332). Nakagawa’s *R*² was 0.053 (marginal) and 0.388 (conditional), indicating that the fixed predictors alone explained 5.3% of the variance, rising to 38.8% once bromeliadL and dateLlevel clustering was accounted for.

On the logLcount scale, the bromeliadLlevel randomLintercept variance was 0.30 (standard deviation [SD] =L0.55) and the samplingLdate variance 0.69 (SDL=L0.83); approximating the levelL1 (residual) variance as one gave a bromeliad intra-class correlation coefficient (ICC_bromeliad_) of 0.15 (15L% of total variance) and ICC_date_ of 0.35 (35L% of total variance), indicating that counts were more strongly clustered by sampling week than by individual plant.

Of the fixed effects (Table 2), patch size was significant: each additional bromeliad was associated with a 4% increase in expected immature□mosquito counts (incidence-rate ratio [IRR]□1.04; 95□%□CI□1.02–1.05; *P*□<□0.001; Figure 5). Bromeliad species *A. imperialis* plants contained 31% more immature mosquitoes than *A. nigripetala* (IRR□1.31; 95□%□CI□0.97–1.77; *P*□=□0.08; Figure 5). The precision around samples from *V. hieroglyphica* was very wide (IRR□1.61; 95□%□CI□0.61–4.28; *P*□=□0.34).

**Figure 5.**
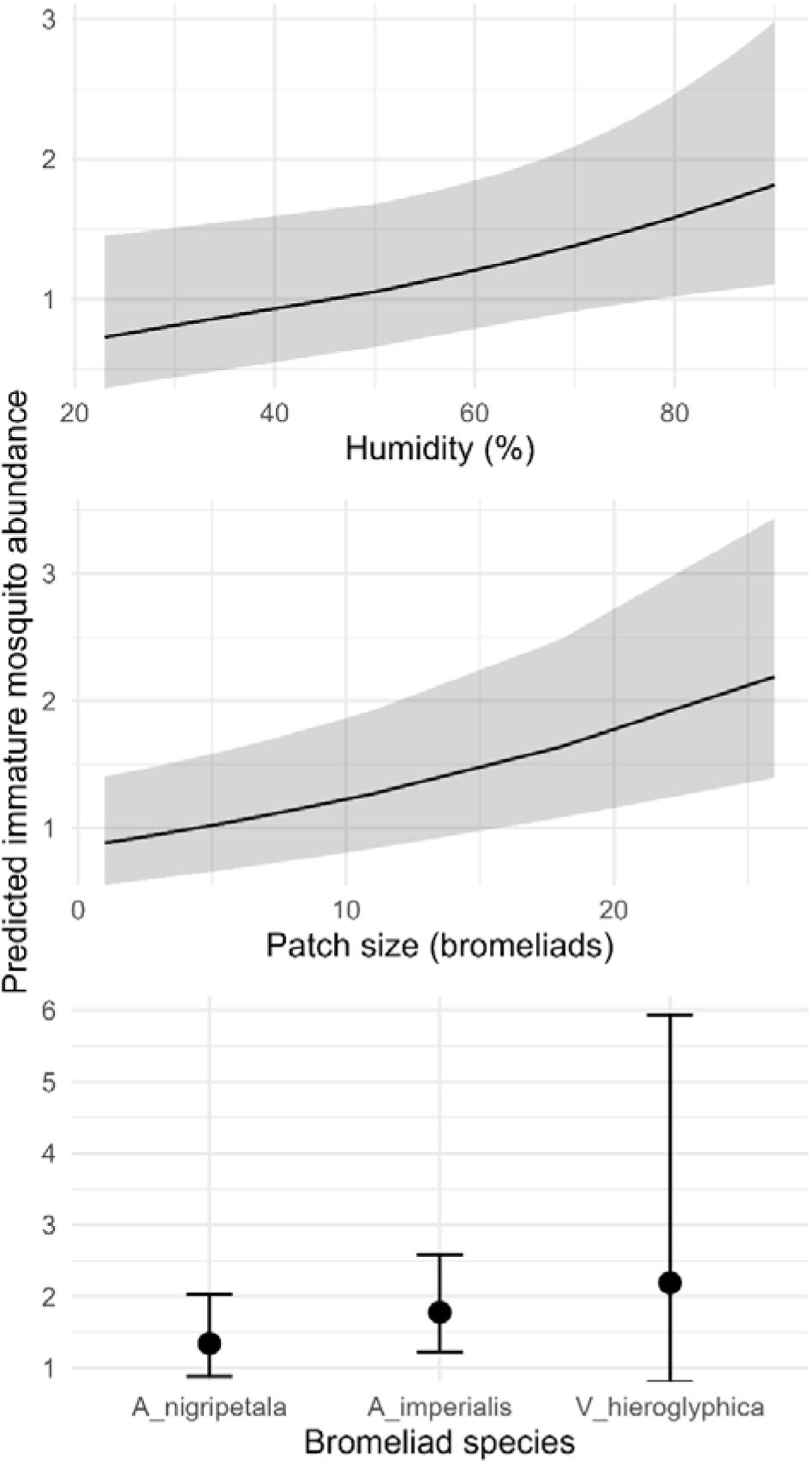
Bias-corrected, population-level predictions from the final negative-binomial GLMM, plotting how (Top) lagged humidity and (Bottom) patch size influence immature-mosquito abundance when all other covariates are held at their mean values (patch size = 6.1 bromeliad plants; 1-week lagged humidity 67.7%, mean temperature = 24.5 °C; total rainfall = 24.9 mm; mean windspeed = 23.6 km/h).

Weekly-lagged humidity showed a positive association (IRR□1.01 per 1□% increase; 95□%□CI□1.00–1.03; *P*□=□0.03). Temperature, rainfall and wind speed effects were non□significant (all *P*□>□0.2; Figure S2)

The multiplicative effect of increased size of bromeliad patch is shown in Figure 6. At the baseline weather conditions (lagged mean weekly humidity□67.7□%, temperature□24.5□°C, windspeed□23.6□km/h, and total rainfall□24.9□mm), a single□plant patch is predicted to contain approximately 1.1 immature mosquitoes (95□% CI 0.7—1.7), while patches of 10 and 20 bromeliads are predicted to contain 14.1 (9.7—20.6) and 37.3 (25.6—54.1) immature mosquitoes, respectively.

**Figure 6.**
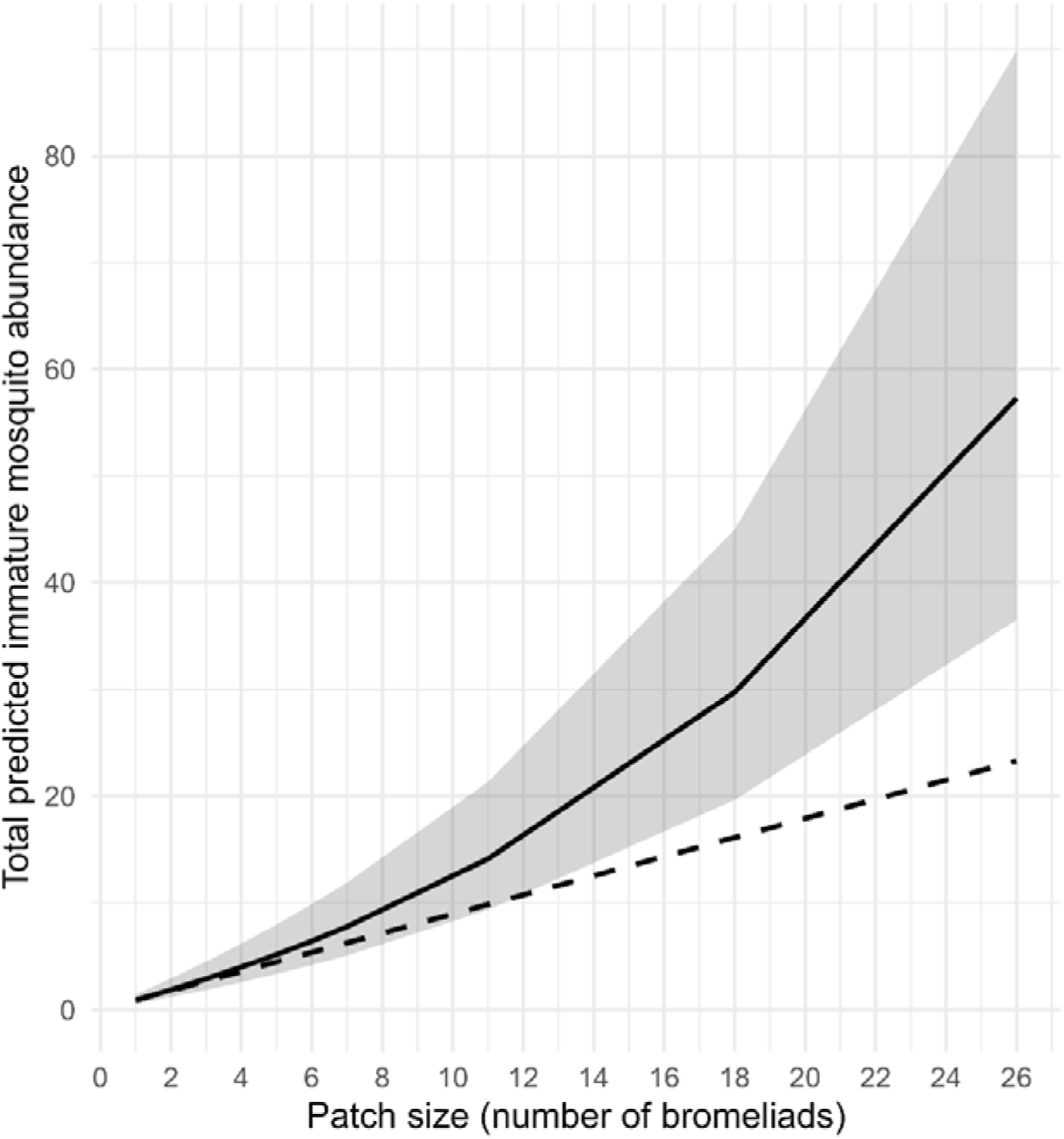
Bias-corrected, population-level prediction of total immature-mosquito abundance as a function of patch size from the final negative-binomial GLMM. Solid line: the model’s multiplicative effect (IRR 1.04 per extra bromeliad). Shaded band: 95% confidence interval. Dashed line: additive baseline (i.e. one-plant rate multiplied by patch size). All other covariates were held at their mean one-week lagged values (humidity = 67.7 %, temperature = 24.5 °C, total rainfall = 24.9 mm, wind speed = 23.6 km/h).

Absolute (additive) effects were quantified via average marginal slopes (FigureLS2). Patch size had the largest additive impact: each additional bromeliad was associated with 0.06 more immature mosquitoes per sampling (95□%□CI□0.02–0.09; *P*□<□0.001). Weekly□lagged humidity showed a marginal effect of 0.02 immature mosquitoes/1□% increase (95□%□CI□–0.003–0.044; *P*□=□0.05). Bromeliad species differences mirrored the multiplicative results: compared to *A.*-*nigripetala*, *A.*-*imperialis* supported an average of 0.41 extra immature mosquitoes/plant (95□%□CI□0 -0.06–0.88; P = 0.08), whereas *V.*-*hieroglyphica* did not differ significantly (0.81; 95□%□CI -1.26–2.9; *P*□=□0.44). Temperature (0.04; 95□%□CI -0.03–0.12; P = 0.23), rainfall (0.001; 95□%□CI -0.02–0.01; P = 0.76) and wind speed (–0.003; 95□%□CI□0.03–0.02; P = 0.80) did not have significant additive effects on immature mosquito counts.

### 3.3 Adult mosquito abundance, species richness, and dynamics

A total of 6,366 adult mosquitoes were collected from 18 traps at six trap sites on the four sampling dates from November 2023 to March 2024 (range 821— 3,221/sampling date, with trap site median). Mosquitoes comprised ten species from four genera (Table 1). The most abundant mosquito identified was *Ae. notoscriptus* (n = 5782), followed by *Cx. quinquefasciatus* (n = 527), and *Cx. annulirostris* (n = 19; Tables 1 and S1). All other mosquitoes comprised < 1% of the total specimen collections (Figure 7). No invasive mosquito species were detected.

**Figure 7.**
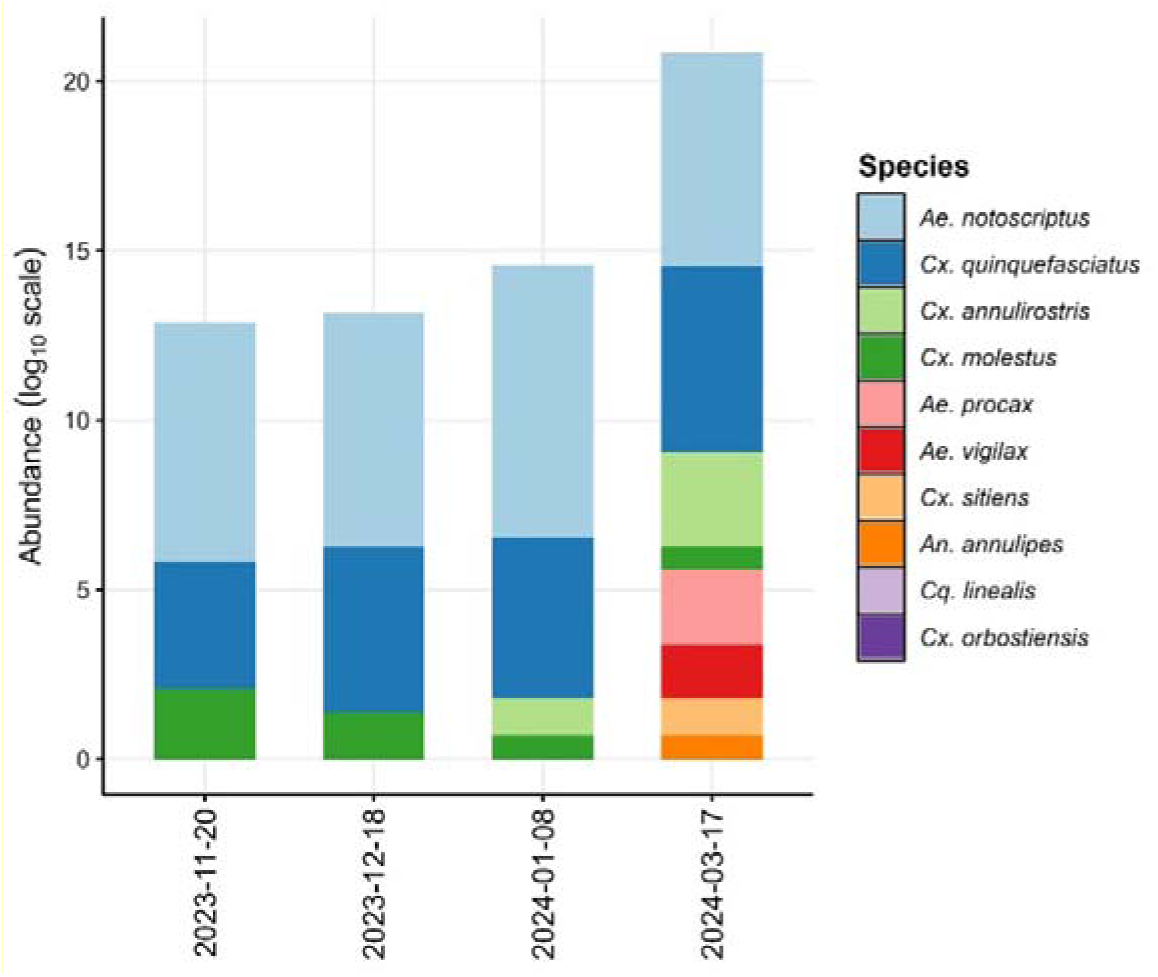
Abundance (log_10_ scale) of adult mosquito species at Camperdown campus, The University of Sydney from November 2023—March 2024.

**Figure 8.**
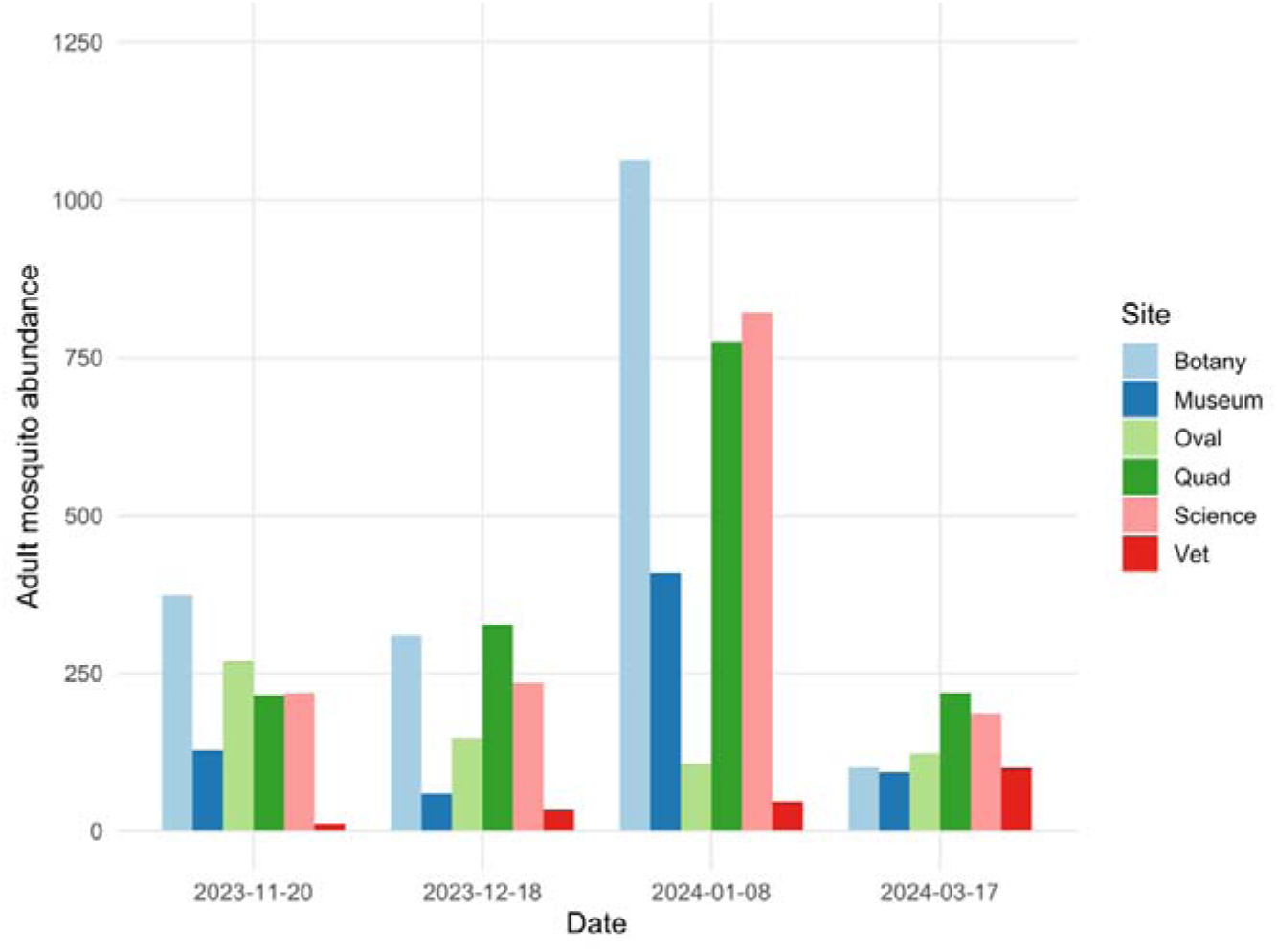
Adult mosquito abundance at six locations at Camperdown campus, The University of Sydney from November 2023—March 2024.

**Table 1.**
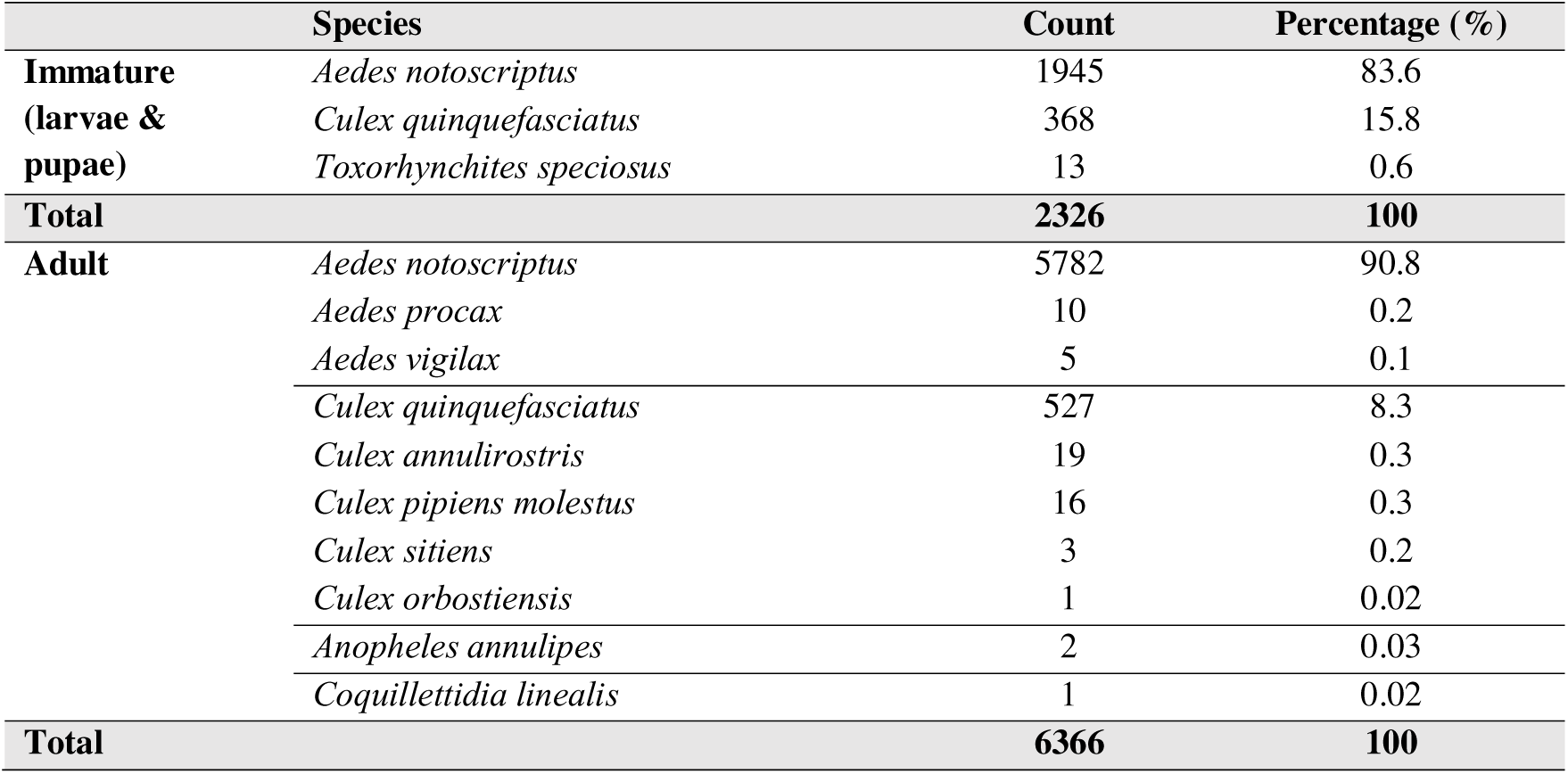
Count of immature and adult mosquito species and their relative abundance at Camperdown campus, The University of Sydney, October 2023—April 2024.

**Table 2:** Fixed-effect estimated incidence rate ratios (IRRs) from the negative-binomial GLMM of immature mosquito abundance in large bromeliads at Camperdown campus, The University of Sydney October 2023—April 2024. CI = confidence interval.

| Variable | IRR | Standard error | Z (Wald) | 95% CI | P value |
| --- | --- | --- | --- | --- | --- |
| Intercept | 0.116 | 0.106 | -2.37 | 0.020–0.689 | 0.02 |
| Temperature (°C) | 1.03 | 0.023 | 1.26 | 0.98–1.07 | 0.21 |
| Humidity (%) | 1.01 | 0.006 | 2.16 | 1.00–1.03 | 0.03 |
| Rainfall (mm) | 1.00 | 0.002 | 0.30 | 0.99–1.00 | 0.76 |
| Windspeed (km/hour) | 0.998 | 0.008 | -0.25 | 0.98–1.01 | 0.89 |
| Bromeliad species:<br>*A. imperialis* | 1.31 | 0.202 | 1.76 | 0.97–1.77 | 0.08 |
| Bromeliad species<br>*V. hieroglyphica* | 1.61 | 0.803 | 0.96 | 0.61–4.28 | 0.34 |
| Patch size | 1.04 | 0.008 | 4.42 | 1.02–1.05 | <0.001 |

**Table 3:** Fixed-effect estimated incidence rate ratios (IRRs) from the negative-binomial GLM of adult mosquito abundance at six sites at Camperdown campus, The University of Sydney October 2023—April 2024. CI = confidence interval.

| Variable | IRR | Standard error | Z (Wald) | 95% CI | P value |
| --- | --- | --- | --- | --- | --- |
| Intercept | 922 | 1303 | 5.25 | 75.6—13018 | <0.001 |
| Temperature (°C) | 0.95 | 0.096 | -5.05 | 0.78—1.16 | 0.61 |
| Rainfall (mm) | 0.97 | 0.009 | -3.31 | 0.95—0.99 | <0.001 |
| Windspeed (km/hour) | 1.07 | 0.125 | 0.54 | 0.85—1.34 | 0.59 |
| Low density | 0.37 | 0.091 | -4.06 | 0.23—0.60 | <0.001 |

Throughout the study period, the mean trap abundance was 88 adult mosquitoes (median; range 0—689). The maximum and minimum abundance from any site was Botany Lawn on 8 January 2024 (n = 1063) and Vet Precinct on 20 November 2023 (n = 11), respectively.

An initial Poisson GLMM with aggregated site counts included bromeliad density (high or low), weather covariates (temperature, rainfall, wind speed; humidity removed due to collinearity), and random intercepts for Site (n = 6) and Date (n = 4) to account for repeated measures. The Date random intercept was dropped because Date variance was estimated as zero and collinear with weather. Equivalent Poisson and negative binomial GLMMs demonstrated a large difference in AICc (986 and 306, respectively). The negative binomial GLMM diagnostics indicated limited dispersion (dispersion = 0.42, *P* = 0.17), no zero inflation (P = 1), and identical marginal (fixed-effects only) and conditional (fixed + random) *R²,* justifying removal of the Site random effect (ICC_Site_ = 0). In the final negative GLM model, Nagelkerke’s pseudo-*R*² was 0.62 indicating that the fixed predictors explained 62% of variance.

Low□density sites had 63% fewer adult mosquitoes than high□density sites, (IRR□=□0.37, 95%□CI□0.23–0.60; P□< 0.001). Each additional millimeter of weekly rainfall was associated with a 3% decrease in expected adult mosquito counts (IRR□=□0.97 (95%□CI□0.95–0.99; P < 0.001). Temperature and wind speed effects were not associated with adult mosquito counts in the model.

## 4 Discussion

This study presents the first systematic evaluation of bromeliads as sources of mosquitoes in Sydney’s urban landscape. Mass-planted, large bromeliads supported significantly higher populations of immature mosquitoes compared to single plantings, and *A. imperialis* could harbour greater numbers of immature mosquitoes compared to other large bromeliad species in this study (*A. nigripetala and V. hieroglyphica*). Sites with greater bromeliad density also yielded higher adult mosquito trap counts. These results add further evidence to the understanding that anthropogenic landscape alteration, including urban plantings, can increase mosquito public health risks (Perrin et al., 2022; Vora, 2008). Recognising bromeliads as productive mosquito habitats has important implications for sustainable urban design and public health management in Australian cities (Mocellin et al., 2009; Wilke et al., 2018).

Beyond expected seasonal variation (NSW Health, 2024), bromeliad patch size was an important predictor of the abundance of immature mosquitoes, with each additional bromeliad plant increasing immature counts by a further 4% per plant across all the other large bromeliads in the patch. This indicates that mass plantings, typical of landscaping practices aiming to maximize green space or cooling capacity (Bartesaghi-Koc et al., 2020; R. M. Karade et al., 2017), are associated with potential exponential increases in immature mosquitoes. This is consistent with findings from Neotropical and subtropical regions, where bromeliad-rich urban gardens have been linked to enhanced mosquito productivity (Frank & Lounibos, 2009; Mocellin et al., 2009; Wilke et al., 2018). Bromeliads alone, or other plantings in urban green space, may not be the sole contributor to local mosquito populations given the presence of other breeding sites or landscape microclimates (Ladino et al., 2019; Medeiros-Sousa et al., 2017), underscoring the need for integrated mosquito habitat management in designed urban ecosystems. However, the selection of bromeliad species and their planting densities, in urban green space warrants consideration with regard to potential mosquito risk.

Studies in the Neotropics and subtropics have reported strong associations between specific bromeliad species’ physical traits – such as water storage capacity, size, and leaf arrangement – and local mosquito diversity and productivity (Cardoso et al., 2015; Navarro et al., 2007). In the present study, *Ae. notoscriptus*, a recognised container-breeding species (Kay et al., 2008; Metzger et al., 2021), was the most abundant mosquito in bromeliads, and the largest species surveyed, *A. imperialis*, may offer both greater oviposition site volume and more stable microclimates than smaller species, conditions likely to promote larval survival (Bentley & Day, 1989; Marino et al., 2011).

Both immature and adult collections were dominated by *Ae. notoscriptus* and *Cx. quinquefasciatus*, consistent with prior Sydney surveys in urban-dominated areas (C. Webb et al., 2001, 2016a). The presence of these species in large numbers where mass bromeliad plantings occur suggests these plants may contribute to local mosquito populations and, potentially, to human arbovirus risk (Jansen et al., 2008; Watson & Kay, 1998). Known to exploit a wide range of container habitats, including bromeliads, these mosquitoes can increase pest and public health risks when conditions favour their proliferation (Allan et al., 2005; Cardoso et al., 2015; Kay et al., 2008). The detection of larvae from the predatory mosquito *Tx. speciosus* also highlights the potential for complex trophic interactions within these phytotelmata (Ceretti-Junior et al., 2015; Frank & Lounibos, 2009), warranting further investigation into predator–prey dynamics and their influence on vector species abundance.

Local weather variables had less influence on immature and adult mosquito abundance than the local contexts of bromeliad abundance, density and species. The positive effect of humidity on immature mosquito counts is consistent with literature linking higher humidity to increased mosquito survival and oviposition (Drakou et al., 2020; Khan et al., 2018), and rainfall could have reduced adult counts due to larval flushing (Tian et al., 2015). Our findings are consistent with previous studies and meta-analyses that have demonstrated that local habitat characteristics are often strong predictors of urban mosquito abundance (Perrin et al., 2022; Roiz et al., 2014). The ecological dynamics within bromeliads, like predator-prey relationships and microclimate, warrant further research as previous investigations of urban mosquito habitats demonstrate the need for understanding site-specific biotic and abiotic factors influencing the activity of mosquitoes (Frank & Lounibos, 2009; Hanford et al., 2020).

It is important to note that the presence of mosquitoes in bromeliads does not necessarily result in public health threats associated with mosquito-borne disease. While many urban areas globally where species such as *Ae. aegypti* or *Ae. albopictus* are present, there will remain a threat of transmission of viruses such as DENV. However, in Australia where the most prevalent mosquito-borne disease are enzootic in nature (e.g. RRV), the presence of key vertebrate reservoir species (for example, native marsupials) will also determine disease risk (Claflin & Webb, 2015). Notwithstanding the risk of endemic mosquito-borne diseases, the use of bromeliads in urban plantings may enhance conditions for exotic container-inhabiting species such as *Ae. albopictus* or *Ae. aegypti* (Nicholson et al., 2014; C. E. Webb et al., 2021).

While this study provides evidence linking high-density bromeliad plantings and mosquito production, several limitations should be noted. Only the central tanks of each large bromeliad were sampled; immature mosquitoes dwelling in peripheral leaf axils might have been missed, potentially underestimating total mosquito productivity as observed in South American surveys (Ceretti-Junior et al., 2015; Frank & Lounibos, 2009). Internal bromeliad factors such as unmeasured predators or competing larvae could have influenced counts in additional to external factors (Lounibos, 2007), and the focus on a single university campus with a limited number of other water bodies or habitats reduces generalisability to urban areas with greater habitat diversity (Medeiros-Sousa et al., 2017). Broader, multi-site, and multi-habitat longitudinal studies, including thorough survey of the diverse urban aquatic habitats, are needed to clarify the full ecological and public health significance of bromeliad-associated mosquito production (Wilke et al., 2018).

## 5. Conclusion

Our findings suggest that mass plantings of large, water-holding bromeliads can significantly increase populations of pest and vector mosquito species in urban environments. Given contemporary moves for sustainable landscaping and urban greening, planners and local authorities should carefully consider the density and placement of bromeliads or substitute these species with non-water-retaining alternatives and use an integrated approach to manage the risk of vector-borne disease. As human populations in urban landscapes grow, proactive management of mosquito habitats in public plantings will become an increasingly important component of public health strategy in Australia and globally.

## CRediT authorship contribution statement

**Pui Yin Lok**: Conceptualization, Methodology, Software, Validation, Formal analysis, Investigation, Writing – Original Draft, Review & Editing, Visualization.

**Victoria Brookes:** Conceptualization, Methodology, Software, Investigation, Resources, Writing – Review & Editing, Supervision.

**Cameron Webb:** Conceptualization, Methodology, Investigation, Resources, Writing – Review & Editing, Supervision.

## Ethical approval and consent to participate

Not applicable

## Consent of publication

Not applicable

## Declaration of interests

The authors declare that they have no known competing financial interests or personal relationships that could have appeared to influence the work reported in this paper.

## Declaration of Generative AI and AI-assisted technologies in the writing process

During the preparation of this work the author(s) used ChatGPT 4.0 in order to revise coding for data analysis in R. After using this tool/service, the author(s) reviewed and edited the content as needed and take(s) full responsibility for the content of the publication.

## Acknowledgement

The authors acknowledge and pay respect to the people of the Eora, Dharug and Tharawal Nations on whose lands this research was carried out and pay respect to their ancient connection to the natural environment: an understanding far older, deeper and more complex than modern science can comprehend. We acknowledge the assistance of Christine Hong, Sydney School of Veterinary Science, The University of Sydney, for assistance with field work and the Information Hub of The University of Sydney in providing suggestion on the data analysis. NSW Health Pathology receives funding from NSW Health.

## Funding

None.

## Notes

### Competing Interest Statement

The authors have declared no competing interest.

