## Supplementary material for "Mass-planted large bromeliads in an urban landscape increase the risk for mosquitoes of pest and public health concern": Figure S1; Figure S2; Figure S3

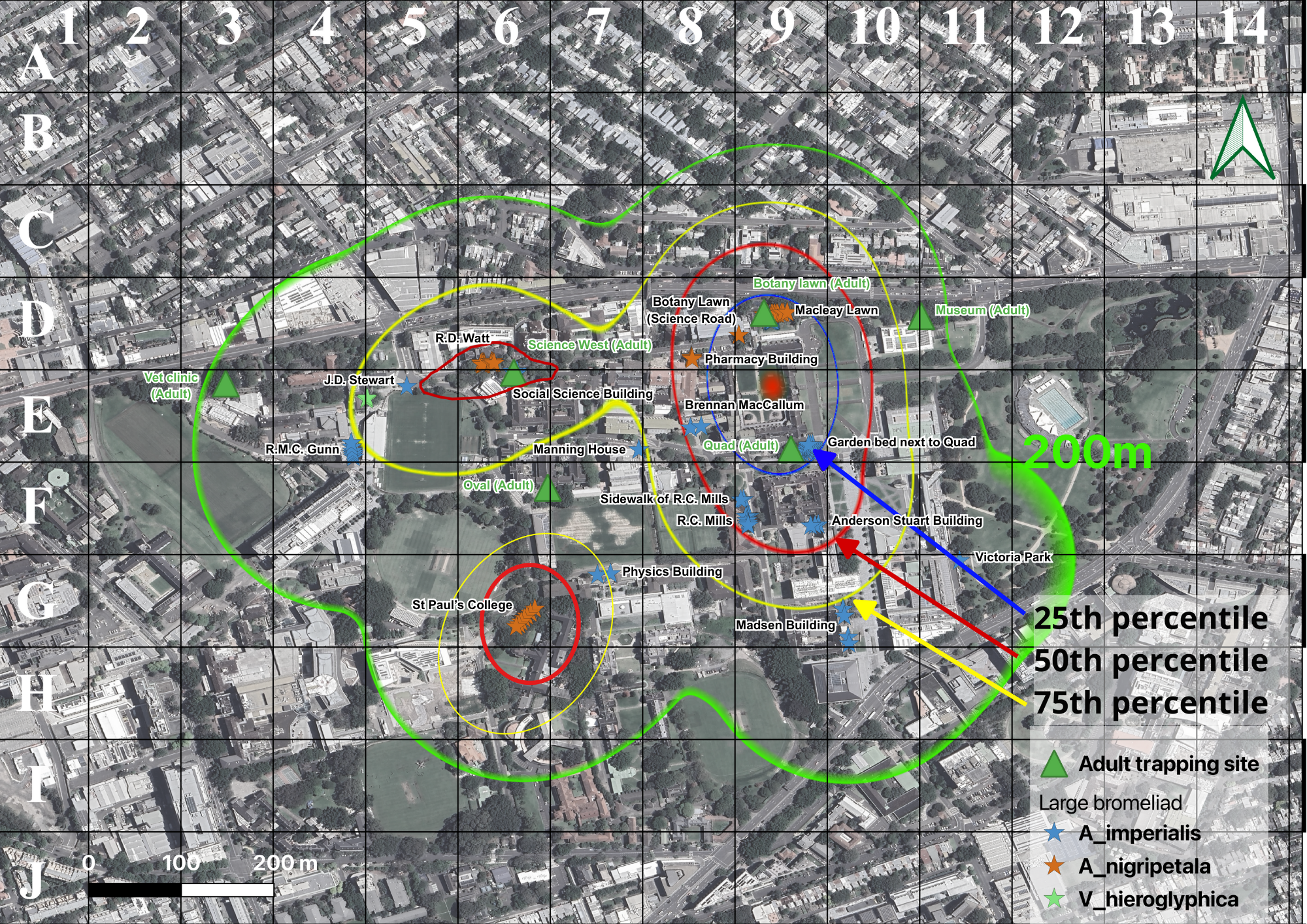

**Figure S1** Study area showing 17 immature and 6 adult mosquito sampling locations in the Camperdown Campus, The University of Sydney to explore the abundance and species richness of mosquitoes between October 2023 and April 2024. QGIS 3.36 was used to create the figure (qgis.org; accessed on 10 October 2023).

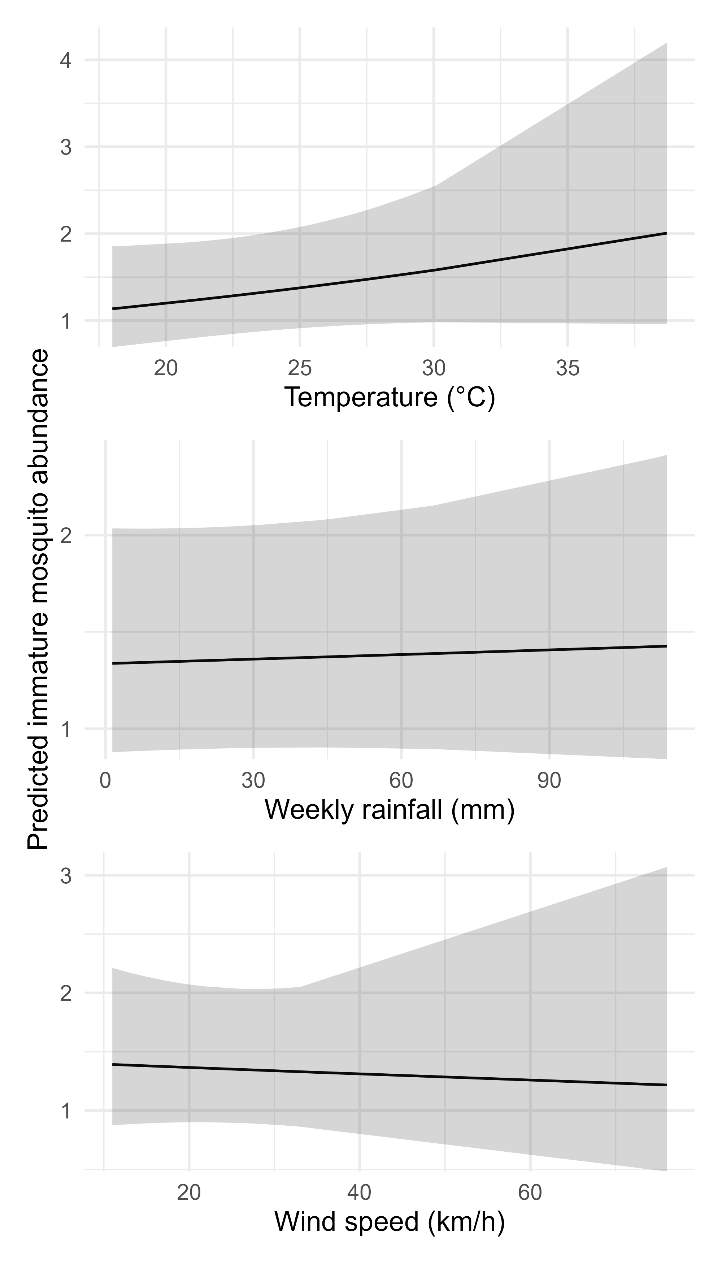

Figure S2: Bias‑corrected, population‑level predictions from the final negative‑binomial GLMM, plotting how 1 week lagged (Top) temperature, (Middle) weekly rainfall, and (Bottom) patch size influence immature‑mosquito abundance when all other covariates are held at their mean values (patch size = 6.1 bromeliad plants; 1 week lagged humidity 67.7%, mean temperature = 24.5 °C; total rainfall = 24.9 mm; mean windspeed = 23.6 km/h).

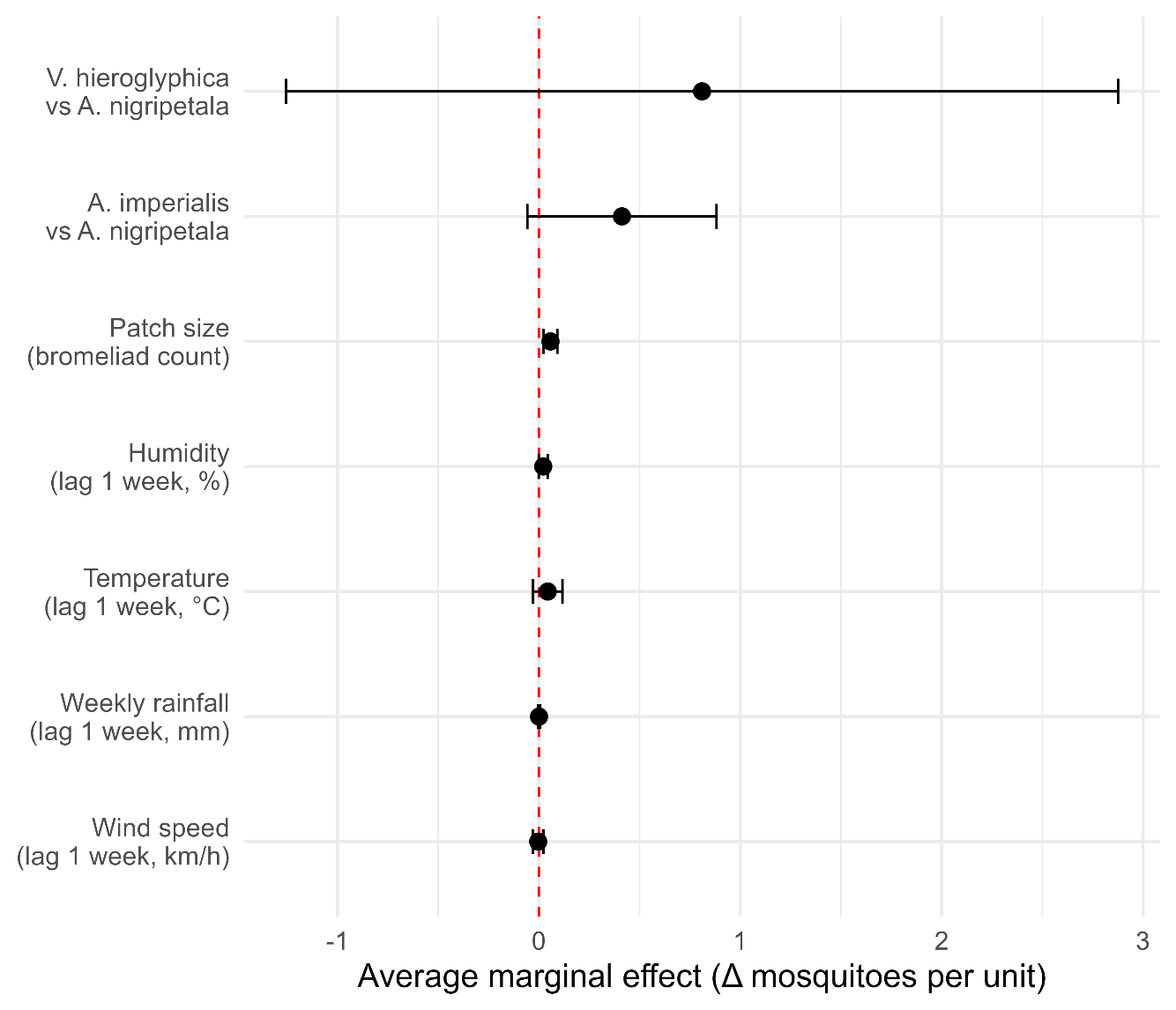

Figure S3: Average marginal effect for fixed effects in the final negative‑binomial GLMM of weather, species, and patch size on immature mosquito abundance in large bromeliads at Camperdown Campus, The University of Sydney, September 2023—April 2024.
